# Stepwise chromatin and transcriptional acquisition of an intraepithelial lymphocyte program

**DOI:** 10.1101/2020.06.04.134650

**Authors:** Mariya London, Angelina M. Bilate, Tiago B. R. Castro, Daniel Mucida

**Author notes:** Contributed equally (Mariya London, Angelina M. Bilate, Tiago B. R. Castro). Correspondence should be addressed to D.M., P., F.

## Abstract

Mesenteric lymph node (mLN) T cells undergo tissue adaptation upon migrating to intestinal lamina propria (LP) and intraepithelial (IE) compartments, ensuring appropriate balance between tolerance and resistance. By combining mouse genetics with single-cell and chromatin analyses, we addressed the molecular imprinting of gut epithelium on T cells. Transcriptionally, conventional and regulatory (Treg) CD4^+^ T cells from mLN, LP and IE segregate based on the gut layer they occupy; trajectory analysis suggests a stepwise loss of CD4-programming and acquisition of an intraepithelial profile. Treg fate–mapping coupled with RNA– and ATAC–sequencing revealed that the Treg program shuts down before an intraepithelial program becomes fully accessible at the epithelium. Ablation of CD4 lineage–defining transcription factor ThPOK results in premature acquisition of an IEL profile by mLN Tregs, partially recapitulating epithelium imprinting. Thus, coordinated replacement of circulating lymphocyte program with site–specific transcriptional and chromatin changes is necessary for tissue imprinting.

## Introduction

The plethora of food and bacteria derived antigens in the intestinal lumen is separated from the lamina propria (LP) by a single layer of epithelial cells, comprising a highly stimulating environment for the vast underlying immune system ^1, 2, 3^. Immune cells in the LP, intraepithelial compartment (IE), as well as in the gut–associated organized lymphoid tissue daily assimilate luminal nutrients and molecules through a controlled balance between tolerance and resistance^4^. A central mechanism for intestinal tolerance is mediated by Foxp3^+^ regulatory T cells (Tregs)^5, 6, 7^, which can be induced by dietary and microbiota antigens in a TGF-β and retinoic acid dependent manner in the gut–draining mesenteric lymph nodes (mLN), and primarily reside in the LP^8, 9, 10, 11^. Likewise, TGF-β and retinoic acid, along with T-bet, are required for the differentiation of CD4^+^ intraepithelial lymphocytes (CD4-IELs) from both conventional CD4^+^ T cells (Tconv) and Tregs upon migration to the IE in a microbiota-dependent manner^9, 11, 12, 13, 14, 15^. CD4-IELs have also been implicated in controlling local inflammation^9, 12^. The hallmarks of CD4-IEL conversion include co-expression of CD4 and the CD8αα homodimer which is induced by a switch from conventional CD4-to-CD8 lineage–defining transcription factors. CD4-IELs downregulate ThPOK (encoded by *Zbtb7b)*, expressed by all conventional mature CD4^+^ T cells and upregulate the long form of Runx3, expressed by mature CD8^+^ T cells^9, 12, 16^. While environmental cues and transcription factors involved in the differentiation of naïve CD4^+^ T cells into gut–adapted T cell subsets have been described in the past decade, the transcriptional and chromatin changes that accompany such tissue–specific imprinting have not been elucidated.

We used genetic fate–mapping and gene ablation mouse models coupled with single-cell RNA–, ATAC– and ChIP–sequencing approaches to uncover the molecular mechanisms of how the intestinal tissue can imprint T-cell fate decisions on migrating mLN CD4^+^ T cells. We found that gut–associated CD4^+^T cell transcriptional profiles largely segregate by tissue location, indicating that upon leaving the gut–draining LNs, migrating cells quickly adapt to either LP or IE compartments; IE–adapted cells followed a stepwise acquisition of an IEL profile through a distinct pre-IEL stage. We specifically followed how the generally stable Treg phenotype is destabilized upon T cell migration to the IE compartment. We found that the Treg program is first downregulated at a pre-IEL stage before a cytotoxic IEL program is made accessible at the chromatin levels and then subsequently transcribed. Finally, we showed that while natural ThPOK downmodulation marks the pre-IEL stage, premature ThPOK loss in Tregs allows for the expression of the IEL profile before the Treg program is fully shut down. Our studies uncovered wide, tissue–specific and stepwise chromatin and transcriptional changes in T cells upon transitioning from tissue–draining LNs to tissue sites and revealed specific roles for lineage–defining transcription factors in driving this process.

## Results

### Intestinal tissue sites imprint a unique program on migrating CD4^+^ T cells

Both peripheral Tregs and Tconv CD4^+^ T cells further differentiate into CD4-IELs upon migration to the intestinal epithelium^9, 12, 16^. To characterize CD4^+^ T cell heterogeneity in the gut along the draining lymph nodes (LNs)-tissue axis, we combined single-cell RNA-Sequencing (scRNA-Seq) with fate-mapping approaches. First, to simultaneously study the transcriptional changes of Tregs and Tconv as they convert to CD4-IELs, we crossed the *Foxp3*^CreER-eGFP^:*Rosa26*^lsl-tdTomato^ (i*Foxp3*^Tomato^) Treg-fate mapping strain^17^ with *Zbtb7b*^GFP^ reporter mice^18^, the latter representing a strategy to follow IEL development using ThPOK loss as a marker^9^. We administered tamoxifen to label Tregs and follow their fates (as Tomato^+^) in three different sites along the LN-tissue axis: gut-draining mesenteric lymph nodes (mLN), lamina propria (LP) and epithelium (IE). We sorted tomato negative (Tom^−^) or positive (Tom^+^) CD4^+^ T cells from the three locations and pooled the cells per tissue at approximately 2:1 (Tom^−^: Tom^+)^ ratio in order to achieve a better representation of current Tregs (Tom^+^GFP^+^), former Tregs (Tom^+^GFP^−^) and non-Tregs (Tom^−^). We performed droplet-based scRNA-Seq using the Chromium 10X (10X Genomics) platform on the three libraries, allowing us to explore the tissue-dependent relative heterogeneities of 6668 cells (**Supplementary Fig. 1a-c**). As expected from our previous studies^12, 16^, CD4^+^ T cell populations in the epithelium showed low levels of ThPOK primarily in the IE, which correlated with the acquisition of CD8α (**Supplementary Fig. 1a-b**).

Cells were positioned based on gene expression similarities by uniform manifold approximation and projection (UMAP)^19^ and assigned to 25 unbiased clusters (0-24, numbered by size order), including a small cluster of contaminating B cells (cluster 24). The vast majority of cells were segregated by tissue, suggesting that different microenvironments, including those within the gut, play a major role in defining the gene expression programs (**Fig. 1a, b**). We defined the clusters based on their top differentially– expressed genes (**Fig. 1a-c, Supplementary Table 1**). Four clusters (11, 16, 19 and 20) contained cells from at least two different locations **(Fig. 1a)**, and we classified them as mixed clusters (MX). Cluster 20 was defined by genes encoding mitochondrial localized proteins (*Atad3a, Cox7b*), snoRNAs regulators (*Nhp2, Gar1*), and genes related to ER stress such as heat shock proteins (*Hspd1, Hspa9*) (**Supplementary Table 1**), and was thus defined as “stressed” cells. Cluster 16 was characterized by expression of genes related to cell cycle, including *Cdc’s, Cdk’s* and *Ccn’s* and was inferred to contain cycling cells. Cells of cluster 19 expressed high levels of integrins (*Itga4, Itgb1, Itgb7*) as well as *Vim* and *Actg1*, suggesting these are motile cells (**Supplementary Table 1, Supplementary Fig. 1d**). The largest mixed tissue cluster, cluster 11, was partly defined by effector Treg genes (*Izumo1r, Pdcd1, Lag3, Icos, Tnfrsf4*) (**Fig. 1c, d, Supplementary Table 1**). The three largest mLN clusters (MN_0,9,10) were defined by naïve markers (*Sell, Ccr7, Klf2*), as expected given its location (**Supplementary Fig. 1e, Supplementary Table 1**). One of the smallest mLN clusters, cluster 17, was defined by expression of *Actb*, whereas cluster 23 consists of multiple genes encoding interferon-induced proteins (Ifits) (**Fig. 1c, Supplementary Table 1)**. In general, cells isolated from mLN expressed more ribosomal protein-encoding genes in comparison to LP and IE cells, possibly due to their more naïve state and thus lower expression of cell state-defining markers (**Fig. 1b**). As a whole, T cells from the LP expressed genes related to activation such as those in the Jun, Dusp, and NF-κb families (**Fig. 1b, c**). Clusters 3 and 21 consisted of cells with relatively higher levels of *tdTomato, Ctla4, Icos*, and other effector Treg genes^20, 21^ (**Fig. 1d, Supplementary Fig. 1f, Supplementary Table 1**). Cells in LP clusters 2 and 15 expressed Th1 (*Il12rb2, Il18r, Stat4*) and Th17 (Il*17a, Il22, Rora*)-associated genes, respectively (**Fig. 1a, c, Supplementary Fig. 1d, Supplementary Table 1**). Clusters 1 and 7 contained cells from LP that did not express the typical Th1 or Th17 signatures but had the effector phenotype. Cluster 7 was polarized towards an IEL-profile, with higher expression of *Nkg7, Gzmk*, and *Ccl5* (**Fig. 1a, c, e, Supplementary Table 1**). We considered all clusters in the IE as *tdTomato*-expressing Tregs (clusters 5,18), *Cd8a*-expressing CD4-IELs (clusters 4,6,13,22) or cells that shared multiple genes with CD4-IELs but lacked *Cd8a expression* (clusters 8, 12, 14) (**Fig. 1a, e, Supplementary Fig. 1d, f-h, Supplementary Table 1**).

**Figure 1.**
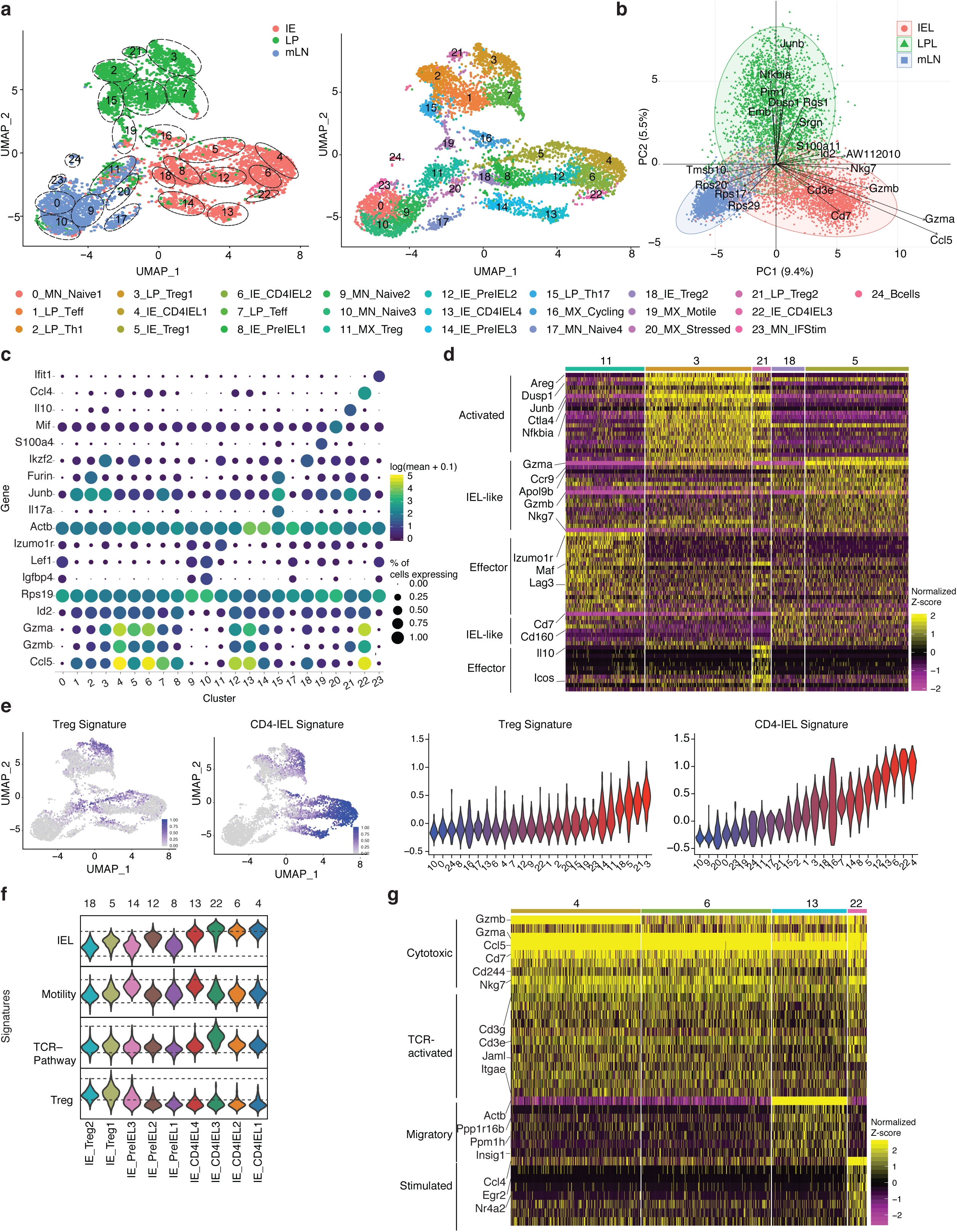
Intestinal epithelium imprints a cytotoxic program on migrating CD4^+^ T cells. **(a-g**) *Foxp3*^eGFP-Cre-ERT2^x*Rosa26*^lsl-tdTomato^x*Zbtb7b*^GFP^(i*Foxp3*^Tom^ThPOK^GFP^) mice were treated with tamoxifen for 10 weeks, Tomato^−^ and Tomato^+^ CD4^+^ T cells from mesenteric lymph nodes (mLN), lamina propria (LP) and intestinal epithelium (IE) were sorted for scRNA-Seq using 10X Genomics platform from one mouse. Sorted Tomato^−^ and Tomato^+^ cells were pooled in a 2:1 ratio per tissue, resulting in 3 separate libraries. (**a**) Uniform manifold approximation and projection (UMAP) representation of all sequenced cells (6668), color coded by tissue of origin with outlined clusters (left) or colored by cluster (right). Cluster names and numbers correspond to colors throughout the figure as indicated. (**b**) Principle component analysis of cells per tissue, exhibiting top variable features. (**c**) Top expressed genes per cluster. Circle size represents proportion of cells per cluster expressing the indicated gene and color represents expression level. (**d**) Heatmap of top differentially expressed genes in Treg clusters represented by the normalized Z-score. (**e**) Expression levels of Treg and CD4-IEL signatures in sequenced cells (left) and per cluster of cells (right). (**f**) Expression levels of indicated signatures among IE clusters as indicated. (**g**) Heatmap of top differentially expressed genes in CD4-IEL clusters by the normalized Z-score.

To gain insight into how the gut environment influences Treg intra-tissue adaptation, we further compared all Treg clusters (11, 3, 21, 18, 5) within the different gut-associated sites (**Fig. 1d, e, Supplementary Table 1**). The two Treg clusters (3,21) present in the LP displayed a more activated profile *(Ctla4, Areg, Dusp1, Junb, Nfkbia)*. Cluster 21 consisted of Tregs expressing *Icos* and *Il10*, while those of cluster 3 expressed higher levels of *Gzma*. Both IE Treg clusters (5, 18) displayed an activated/effector-like phenotype (*Cd3e, Cd3g, Tnfrsf9, Tnfrsf18*), and were also more CD8-like (*Cd160, Cd7*). Additionally, cells in cluster 5 expressed elevated levels of granzymes and *Nkg7*, suggestive of cytotoxic potential. Cells in cluster 11, the only cluster to include mLN Tregs had lower expression of effector Treg markers and of the gut homing receptor *Ccr9* compared to enteric tissue Tregs (**Fig. 1d, e**). These results point to a previously unappreciated level of Treg heterogeneity between closely related, yet distinct tissues, with a skew towards the IEL/cytotoxic program within the epithelium.

We next further characterized the heterogeneity of epithelial CD4^+^ T cells. We defined gene-expression signatures for Tregs, TCR-stimulated cells, motile cells, and CD4-IELs, based on the expression profiles of the different clusters (**Fig. 1e, f, Supplementary Fig. 1d, g**). Although both Treg clusters (5 and 18) expressed the highest levels of Treg-signature genes, cluster 14 also expressed the same genes at a lower level. CD4-IEL cluster 22 displayed the highest level of genes in the TCR pathway (**Fig. 1f, Supplementary Fig. 1g**). Clusters 13 and 14, positioned next to each other, expressed the highest levels of motility-related genes, suggesting increased migratory capacity. Finally, while three out of the four CD4-IEL clusters (4, 6 and 22) displayed equally high CD4-IEL signatures, cluster 13 was slightly less polarized towards the CD4-IEL phenotype (**Fig. 1 e-g, Supplementary Fig. 1d, g**). The three non-CD4-IEL and non-Treg clusters in the IE (8, 12, 14) nevertheless displayed a high level of the CD4-IEL signature (**Fig. 1e**). These clusters were characterized by the presence of cells expressing equal or slightly lower levels of IEL markers such as *Itgae, Cd160, Nkg7* compared to *bona fide* IELs (**Supplementary Fig. 1g, h, Supplementary Table 1**), suggesting that these constitute a pre-IEL population. Among these pre-IEL clusters, which were very similar to each other, cluster 12 was the most similar to CD4-IELs expressing higher levels of *Ccl5* and *Gzmb* (**Fig. 1e, f, Supplementary Fig. 1g, h**). Likewise, the CD4-IEL clusters were separated by only minor differences, including upregulation of genes in the TCR pathway in cluster 22 (*Egr2* and *Nr4a2*) and of a motility signature in cluster 13 (*Actb* and *Ppp1r16b*). All CD4-IELs expressed *Cd8a, Cd244, Gzma, Cd7, Nkg7, Jaml* as well as *Ccl5* **(Fig. 1g, Supplementary Table 1**). Of note, tomato-positive and -negative CD4-IELs were dispersed among all CD4-IEL clusters, indicating that CD4-IELs derived from Tregs or from Tconvs did not display major differences capable of overriding their CD4-IEL signature similarities (**Supplementary Fig. 1f**). Together, our data indicate that CD4^+^ T cells in the IE are placed on a gradient leading to the full expression of a CD4-IEL signature.

To understand how the gradient of transcriptional changes that culminates in tissue adaptation of CD4^+^ T cells within the lymphoid to non-lymphoid tissue axis, we performed pseudotime analysis that produce cell trajectory inferences using two independent but complementary methods, Monocle3 and Slingshot (**Fig. 2a**). While Monocle3 constructs a minimum spanning tree utilizing the individual cells and projects it onto the UMAP embedding, Slingshot uses the cluster centers making it less sensitive to outliers. Color– coding cells along differentiation paths showed that cells making up the clusters along the edges of the LP and IE compartments in the UMAP, including all CD4-IEL clusters, are the most “terminally– differentiated” compared to mLN cells. (**Fig. 2a**). The trajectory branches leading to LP versus IE clusters bifurcated within the mixed-tissue cluster 11 and no further connections were detected between LP and IE (**Fig. 2a**), suggesting that once cells acquire a LP profile they likely remain there. In agreement with the UMAP, pre-IELs were positioned as intermediate stages in the trajectories from naïve CD4^+^ T cells of the mLN to CD4-IELs (**Fig. 2a**). Based on the gradient towards the IEL profile within the epithelium as well as trajectory analysis, we chose three combinations of clusters along the mLN-IE axis, and ordered their cells based on principal component analysis (PC1). The diagonal distribution of cells in all combinations corroborated the differentiation pattern from least differentiated mLN cells to terminally differentiated CD4-IELs (**Fig. 2b**). This progression consisted of the gradual upregulation of genes such as *Ccl5*, granzymes, *Nkg7 and Cd8a*, until their terminal IEL differentiation is complete (**Fig. 2b, c**). These genes were also the main drivers of the IE-signature described in figure 1, further suggesting that the IEL profile defines epithelial imprinting. Our scRNA-Seq revealed that while the lamina propria allows for the maintenance of distinct CD4^+^ T cell subsets, this heterogeneity is not seen in the epithelium, suggesting that localization to the epithelium is responsible for acquisition of the IEL profile.

**Figure 2.**
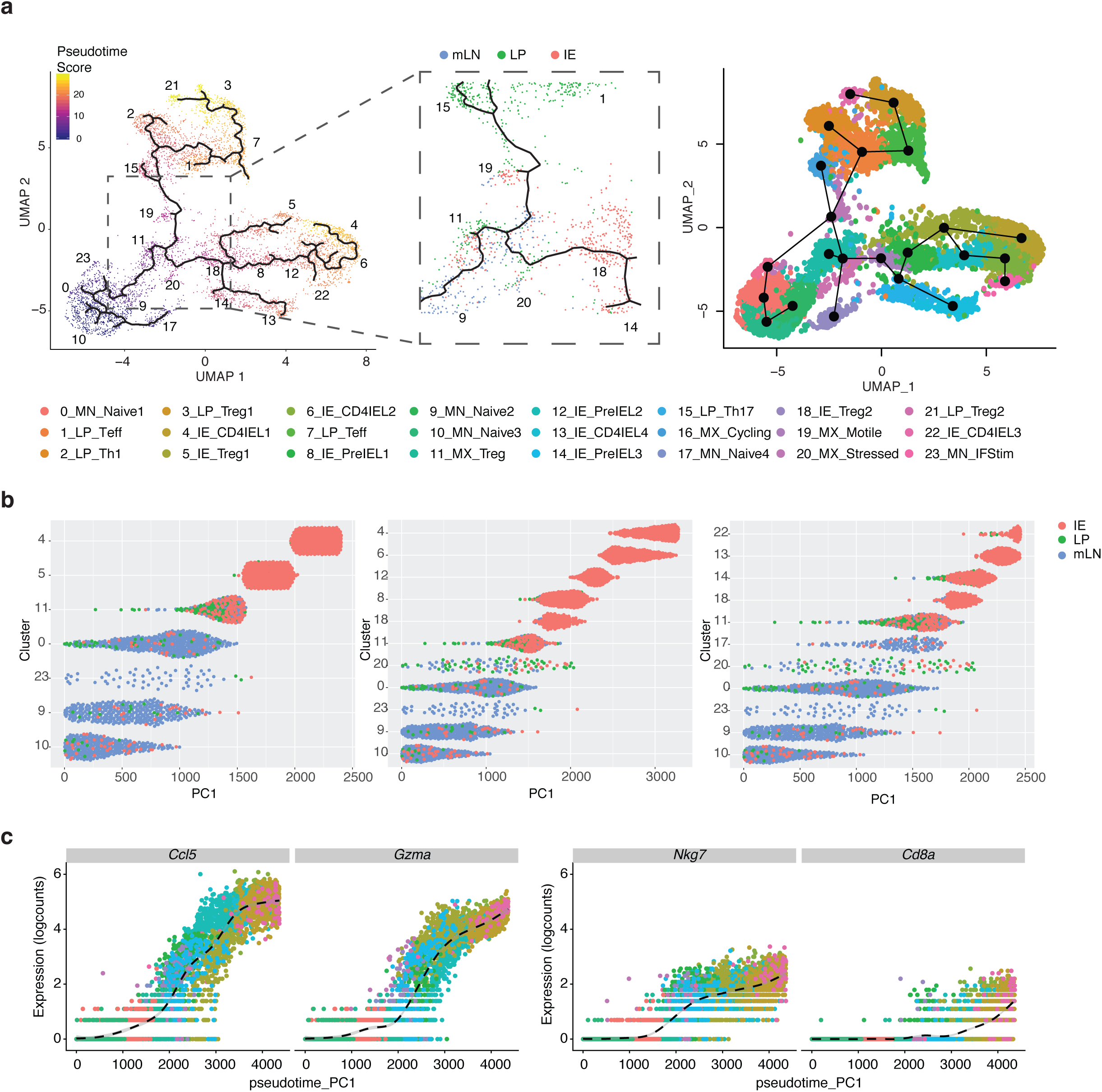
Tissue segregation precedes acquisition of an IEL profile. **(a-c**) *Foxp3*^eGFP-Cre-ERT2^x*Rosa26*^lsl-tdTomato^x*Zbtb7b*^GFP^(i*Foxp3*^Tom^ThPOK^GFP^) mice were treated with tamoxifen for 10 weeks, Tomato^−^ and Tomato^+^ CD4^+^ T cells from mesenteric lymph nodes (mLN), lamina propria (LP) and intestinal epithelium (IE) were sorted for scRNA-Seq using 10X Genomics platform. Sorted Tomato^−^ and Tomato^+^ cells were pooled in a 2:1 ratio per tissue, resulting in 3 separate libraries. (**a**) Pseudotime analysis of sequenced cells using Monocle3 with cells color-coded by pseudotime gradient (left) and by Slingshot with cells color-coded by cluster (right). Numbers indicate the UMAP clusters. Middle panel displays a point of bifurcation in the Monocle3 trajectory analysis, cells colored by tissue. Numbers indicate the UMAP clusters. (**b**) Cells ordered according to principle component 1 (PC1) in. (**c**) Scatter plot of gene expression ranked based on PC1.

### Coordinated transcriptional and chromatin changes during T cell adaptation to the epithelium

We next sought to couple the stepwise changes in transcription with those of chromatin accessibility as Tregs destabilized to become CD4-IELs. As noted in previous studies^12, 16^, ThPOK downmodulation precedes CD8α-acquisition. Our scRNA-Seq analysis using the ThPOK^GFP^ reporter mouse revealed intermediate stages within the IE, and thus we refer to ThPOK^low^CD8α^−^ cells in the following studies as pre-IELs. We performed assay for transposase-accessible chromatin followed by sequencing (ATAC-Seq) and bulk RNA-Seq on peripherally induced Tregs (iTreg; neuropilin1^low^Tom^+^GFP^high^CD8α^−^), pre-IELs coming from Tregs (Tom^+^GFP^low^ CD8α^−^), Treg-derived CD4-IELs (exTreg-IEL; Tom^+^GFP^low^ CD8α^+^) and Tconv-derived CD4-IELs (Tom^−^GFP^low^CD8α^+^) sorted from the IE of i*Foxp3*^Tomato^ThPOK^GFP^ mice (**Supplementary Fig. 2a, b**). Although the Tom^−^ CD4-IELs in our system are not derived from Tom^+^ Tregs, we use them as a phenotypic endpoint to define CD4-IELs, and to compare any differences associated with Treg-versus Tconv-derived IELs.

In agreement with previous ATAC-Seq data^22^, the majority of accessible chromatin peaks occurred close to transcriptional start sites (TSS) (**Supplementary Fig. 2c**). All populations revealed a similar profile of chromatin accessibility relative to genome regions, with the highest frequency of accessibility at promoters (**Supplementary Fig. 2d**). However, out of the genes with differentially–accessible chromatin regions (DACR) between the populations surveyed, the majority falls within gene bodies, and not promoter regions, suggesting non-random chromatin changes (**Supplementary Fig. 2e**). The likelihood ratio test (LRT) showed that DACRs were divided into 6 clusters; regions in clusters 1-4 gradually decreased in accessibility, while those in clusters 5-6 become more accessible as Tregs differentiate into CD4-IELs (**Fig. 3a**). Both activation (*Cd28*, Dusp and Nf-κb families) and naïve-related genes (*Sell, Klf2, Ccr7, Il7r*) contained chromatin regions which decrease in accessibility as Tregs destabilized, suggesting a shift in the activation state during cell-type progression. DACRs in this progression also included *Ifng* and *Stat4*, suggesting a role for interferon regulation in the Treg to IEL progression, as previously suggested^13^. As expected, classic Treg (Foxp3, *Il10, Cd83, Il2ra, Areg)* and effector (*Lag3, Pdcd1, Ctla4, Il10, Icos*) genes decreased in accessibility as Tregs develop into IELs. Of note, while most Treg markers appeared in multiple clusters (1-4), *Foxp3* was detected only in the 4^th^ cluster of regions that decreased in accessibility as pre-IELs transitioned to the IEL phenotype. This suggests that regulation of other Treg-signature genes occurs before *Foxp3*, in agreement with previous reports showing that *Foxp3* expression does not always correlate with Treg profile and function^20^. Chromatin regions in clusters 5 and 6, which began to become more accessible at the pre-IEL stage, and peaked in accessibility in IELs, included typical CD8-IEL (*Runx1/2/3, Gzma, Litaf, Fasl)* and homing (*Ccl5, Itga4)* genes. Cluster 5 contained regions that remained equally accessible in IELs derived from Tregs (exTreg-IEL) and Tconv (CD4-IEL), providing further evidence suggesting that the IEL signature overrides previous lineage or stage characteristics. Chromatin regions in cluster 6, which become more accessible at the exTreg-IEL stage, were more open in CD4-IELs. In addition to regions in typical IEL genes, cluster 6 included *Cd8a*, suggesting that it is not until the IEL program is made accessible that *Cd8a* is upregulated (**Fig. 3a, Supplementary Table 2**). Moreover, Wald pairwise comparisons (**Fig. 3b, c, Supplementary Table 3**) as well as Euclidean distance analysis (**Supplementary Fig. 2f**) showed that the largest changes in chromatin accessibility occurred as iTregs destabilized towards the ThPOK-low pre-IEL stage. The majority of the DACRs in this step fell on gene bodies (**Fig. 3b**). A relatively minor number of regions, including that of *Cd8a*, became more accessible as pre-IELs transitioned into CD8α-expressing exTreg-IELs, which were the most similar to CD4-IELs; however, the former did retain a number of accessible regions at Treg genes, including *Foxp3*, when compared to CD4-IELs, suggesting that the chromatin landscape of Treg genes is not completely lost upon transition to the IEL phenotype (**Fig. 3c**). Overall, it is not until the Treg program is set to become less accessible that the chromatin of the IEL program becomes more accessible, suggesting that ThPOK loss at the pre-IEL stage, or the events that lead to it, initiate the Treg plasticity observed at the intestinal epithelium.

**Figure 3:**
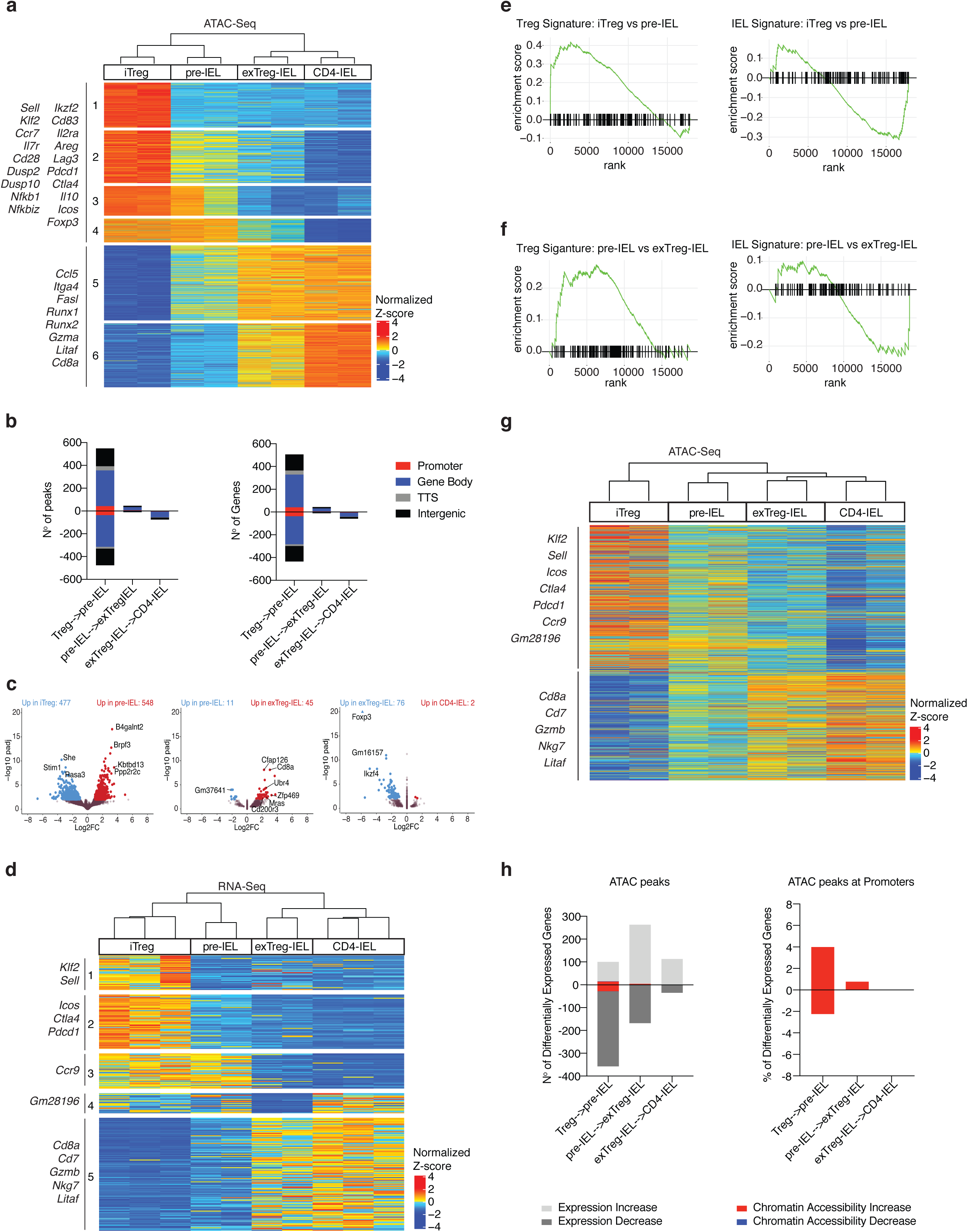
Treg program shutdown precedes IEL programming through a pre-IEL stage. (**a-g**) *Foxp3*^eGFP-Cre-ERT2^x*Rosa26*^lsl-tdTomato^x*Zbtb7b*^GFP^(i*Foxp3*^Tom^ThPOK^GFP^) mice were treated with tamoxifen for 10 weeks and induced Tregs (iTreg; CD4^+^ Tomato^+^ GFP^High^ neuropilin-1^−^ CD8α^−^), pre-IELs (CD4^+^ Tomato^+^ GFP^Low^ CD8α^−^), exTreg-IELs (CD4^+^ Tomato^+^ GFP^Low^ CD8α^+^), and CD4-IELs (CD4^+^ Tomato^−^ GFP^Low^ CD8α^+^) were sorted in bulk from the IE. Assay for transposase-accessible chromatin (ATAC) or RNA libraries were prepared followed by sequencing of indicated populations. (**a**) Heatmap of likelihood ratio test (LRT) of all differentially accessible chromatin regions (DACR) of indicated bulk populations. (**b**) Number of DACR (left) and the genes on which they are positioned on (right) between cell types in sequential progression as performed by Wald pairwise test. Colored by chromosome region as follows: 5’UTR and promoters (Promoter; red), 3’UTR with exons and introns (Gene Body; blue), transcriptional termination site (TTS; grey) and intergenic (black). (**c**) Volcano plots representing DACR between cell types in sequential progression as performed by Wald pairwise test. (**d**) Heatmap of LRT of differentially expressed genes (DEG) between indicated populations. (**e, f**) Gene set enrichment analysis of Treg and CD4-IEL signatures from scRNA-Seq (top panels, cluster 21 and bottom panels, cluster 6, respectively as shown in *Figure 1*) in iTreg to pre-IEL (**e**) and pre-IEL to exTreg-IEL (**f**) progressions. (**g**) Levels of chromatin accessibility among DEG, in the same order as in *d*. (**h**) Numbers of DEGs (left) (light grey for increase, dark grey for decrease) and percent of DACR at promoters among DEG (right) in between cell types in sequential progression as performed by Wald pairwise test as indicated. DACR within those genes (red for increase, blue for decrease in accessibility). Significant DACR p<0.01 and significant DEG p<0.05 in RNA-Seq. Each sample for ATAC-Seq consisted of 5,000-40,000 cells from 6 or 9 pooled mice, n=2 samples. Each sample for RNA-Seq consisted of 300-800 cells per mouse, n=2-3 mice.

To define how changes in chromatin accessibility relate to differentially expressed genes (DEG) in IEL populations, we performed bulk RNA-Seq on the same IE subsets. We also sequenced mLN iTregs in order to compare their transcriptional profile to that of iTregs in the IE (**Supplementary Fig. 2g**). Comparing mLN to IE iTregs confirmed that transcriptional differences segregated by tissue, the latter expressing more activation, intestinal and CD8–associated genes (*Runx3, Cd160, Cbfb, Itgae)*, whereas the former expressed genes associated with resting LN Tregs (*Ccr7, Sell, Klf2*) (**Supplementary Fig. 2g, Supplementary Table 4**). DEGs between all sequenced IE populations revealed 5 unbiased clusters, highlighting the changes in transcriptional profile as Tregs convert to IELs (**Fig. 3d**). At the pre-IEL stage when ThPOK is downmodulated, expression of effector Treg genes (Cluster2; *Icos, Ctla4, and Pdcd1)* was decreased. The clusters of genes down-regulated at the exTreg-IEL stage (Clusters 3 and 4) included multiple pseudogenes and lincRNAs, which could be implicated in cell maintenance and identity integrity, serving as a hallmark of cell differentiation at the exTreg-IEL stage. Finally, cluster 5, which is upregulated by both Treg- and Tconv-derived IELs (exTreg-IELs and CD4-IELs), includes many CD8– and IEL–associated genes (*Cd8a, Cd7, Gzmb, Nkg7, Litaf*). Although these genes begin to be slightly upregulated in the pre-IEL stage, they reach their full levels only when T cells become IELs, as marked by CD8α protein and gene expression (**Fig. 3d, Supplementary Fig. 2a, Supplementary Table 5**). Whereas the signature of cluster 3 maps to the Treg clusters in all tissues in the scRNA-Seq data set, that of cluster 5 is primarily expressed by IE cells, particularly CD4-IELs (**Supplementary Fig. 2h**). Although there is very low expression of this signature by a few LP cells, this CD4-IEL program is gained in the IE, after cells have left the LP, again suggesting that they are not polarized towards this program prior to entering the IE compartment. Gene set enrichment analysis (GSEA) using Treg and IEL signatures from our scRNA-Seq (Clusters 21 and 6, respectively) revealed that expression of the Treg signature decreases as the IEL signature increases and cells progress from the Treg to pre-IEL stage (**Fig. 3e**). In contrast, the pre-IEL to exTreg-IEL progression displayed only minimal changes in the Treg signature (**Fig. 3f**), since Treg-derived pre-IELs no longer expressed most of the *bona-fide* Treg genes. Overall, the gene expression changes during IEL differentiation suggest that the IEL program begins during the pre-IEL stage, before the full Treg program is shut down but after the downregulation of ThPOK.

To further understand how changes in gene transcription correlate with changes in chromatin accessibility, we plotted the levels of chromatin accessibility of genes that changed in transcription in the same order of the heatmap representing RNA-Seq (**Fig. 3d**). We found that in general, the transcriptional profile correlated with chromatin accessibility of those genes in iTregs and CD4-IELs (**Fig. 3g**). Genes that were transcriptionally downregulated after the iTreg stage remained slightly more accessible in the pre-IEL stage and their chromatin slowly closes as they become IELs. On the other hand, genes that were upregulated at the IEL stage were more accessible already at the pre-IEL stage. This data further consolidates pre-IELs as a *bona-fide* transitional stage between Tregs and exTreg-IELs. Moreover, in each Wald pairwise comparison, we analyzed how chromatin accessibility changed in genes that increased or decreased in transcription levels (**Fig. 3h**). We found that concomitant changes in transcription and chromatin accessibility occurred primarily at the iTreg-to-pre-IEL stage, only with about 6% of changes occurring at promoter regions (**Fig. 3h**.) Transcriptional changes in the following stages were largely not accompanied by changes in chromatin accessibility. Comparison of differentially– accessible chromatin and gene expression as Tregs become IELs indicates that the Treg program begins to shut down and IEL program is initiated only at the pre-IEL stage. The transitioning pre-IEL stage is not only marked by ThPOK loss, but also by a transcriptional shut down of the Treg program, which marks the start of chromatin accessibility of the IEL program. It is not until CD8α is expressed at the exTreg-IEL stage that the IEL-genes are concomitantly transcribed. Once Tregs fully transition to IELs, they display a very similar transcriptional profile to that of Tconv–derived CD4-IELs, although the former retains a small level of accessible chromatin of the Treg program. In conclusion, once CD4^+^ T cells reach the epithelium, ThPOK is inevitably downmodulated followed by the commencement of the IEL program.

### ThPOK downmodulation together with the epithelial environment is required for IEL differentiation

Our characterization using single–-cell and bulk approaches strongly indicated that ThPOK downmodulation occurs at the crucial step of Treg destabilization to the pre-IEL stage. To mechanistically understand how ThPOK modulation allows for Tregs to proceed to the pre-IEL stage, we assessed possible genes modulated by ThPOK binding using ThPOK chromatin immunoprecipitation followed by sequencing (ChIP-Seq). Due to antibody and cell number limitations for this assay, we performed ThPOK ChIP-Seq on *in vivo* expanded splenic Tregs. We found that the mouse ThPOK binding motif was similar to the published human motif^23^, and the protein preferentially bound to promoter regions, followed by introns (**Fig. 4a, Supplementary Table 6**).

**Figure 4:**
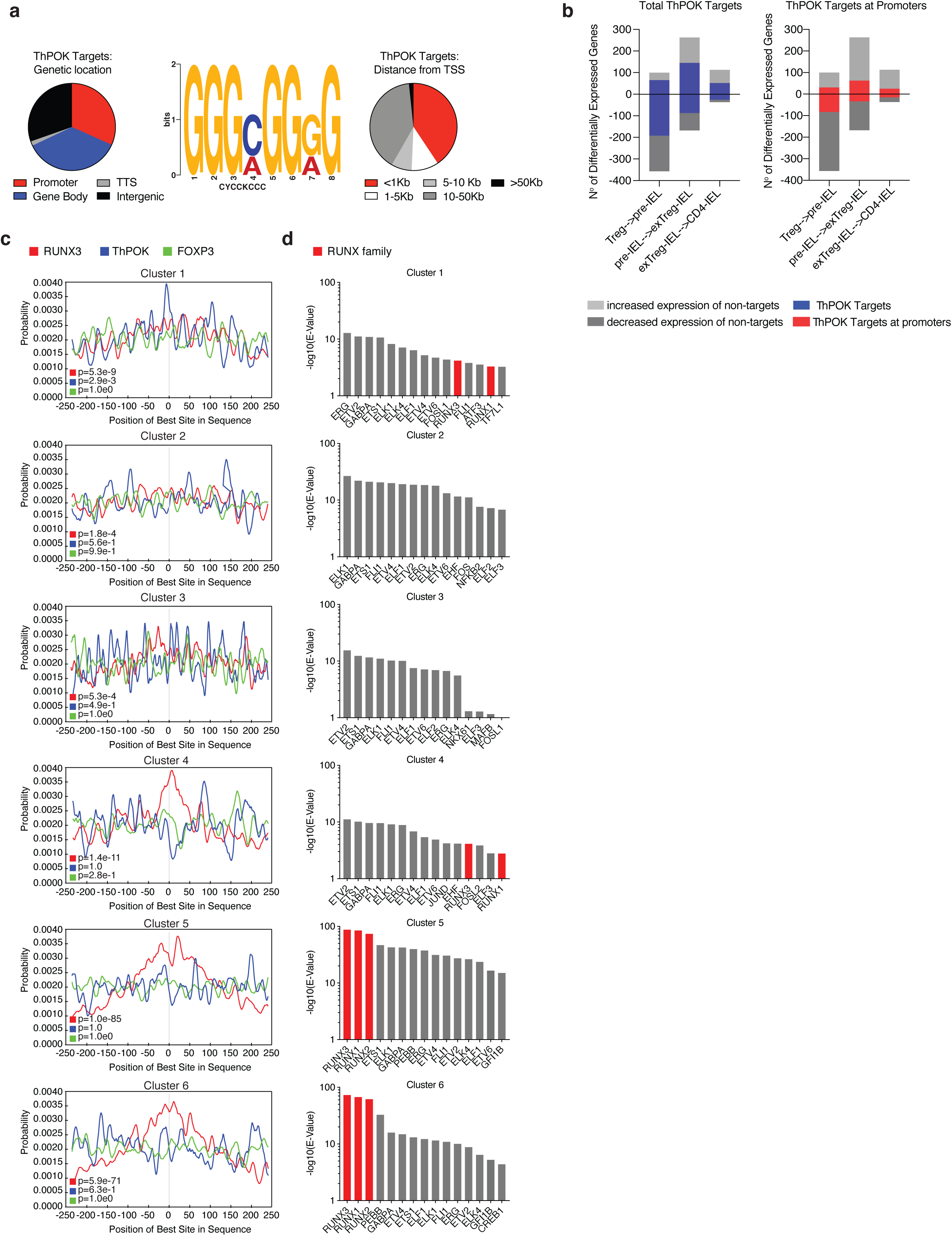
Runx binding motifs are increasingly accessible during the iTreg to IEL progression. (**a-d**) ThPOK ChIP-Seq analysis of *in vivo*–expanded splenic Tregs from *Foxp3*^RFP^ mice coupled with RNA- and ATAC-Seq of iTregs (CD4^+^ Tomato^+^ GFP^High^ neuropilin-1^−^CD8α^−^), pre-IELs (CD4^+^ Tomato^+^ GFP^Low^ CD8α^−^), exTreg-IELs (CD4^+^ Tomato^+^ GFP^Low^ CD8α^+^), and CD4-IELs (CD4^+^ Tomato^−^ GFP^Low^ CD8α^+^) from the IE of i*Foxp3*^Tom^ThPOK^GFP^ mice after tamoxifen treatment (as in *Figures 1, 2*). (**a**) Frequency of ChIP peaks at indicated regions (left) and at indicated distances from transcriptional start sites TSS (right) with *de novo* ThPOK binding motif determined by MEME-ChIP (middle). (**b**) Numbers of differentially expressed genes (light grey for increase, dark grey for decrease) with total putative ThPOK targets (blue; left) or ThPOK targets at promoters (red, right). (**c**) Transcription factor (TF) motif analysis of indicated TFs, collapsed around transcriptional start site (TSS). (**d**) TF motif analysis per cluster of ATAC-Seq LRT heatmap of Figure 3a, performed by MEME-ChIP. Runx family TF’s indicated in red. Significant differentially accessible regions p_adj_<0.01 in ATAC-Seq and significant differently expressed genes p_adj_<0.05 in RNA-Seq. 20×10^6^ cells were used for ChIP-Seq (10% for input, 90% for ThPOK-binding). E-value=1.6e-142 for the ChIP-Seq ThPOK motif.

Overall, genes with ThPOK binding sites were dispersed throughout the Treg to IEL progression **(Fig. 4b**). We assessed the incidence of Runx3, ThPOK (from our analysis) and Foxp3 binding motifs in differentially–accessible chromatin regions during the iTreg to IEL progression. We separately performed this analysis on DACRs from each cluster of the ATAC-Seq heatmap. We found that the Runx3 binding motif was centrally enriched in all clusters, with a marked increase during the acquisition of the IEL program (clusters 5 and 6, **Fig. 4c**). While the ThPOK binding motif was centrally enriched in differentially–accessible chromatin regions only ofin cluster 1, it was accessible in other clusters, suggesting that ThPOK may play a role in both the expression of Treg and repression of IEL genes, as suggested by previous studies targeting ThPOK in mature CD4^+^ T cells^9, 12, 24^. The Foxp3 binding motif was not centrally enriched in any cluster, implying that it may no longer play a functional role in transitioning epithelial Tregs (**Fig. 4c**). Furthermore, to broadly investigate other possible TF regulation that may be correlated with the changes in chromatin accessibility during T cell adaptation to the IE environment, we performed TF motif analysis using MEME-ChIP^25^ on each cluster of the ATAC data (**Fig. 3a, 4d**). Out of the top 15 significantly enriched TF motifs per cluster, those bound by the RUNX family stood out. Accessible chromatin regions with the RUNX binding motif increased during the iTreg to IEL progression, reaching peak significance in clusters 5 and 6. ThPOK downmodulation may leave space for other TF to bind in adjacent DNA regions, a possibility analogous to previously described roles of Runx3 to the CD8-lineage^26, 27^. For instance, Runx3 can bind to ThPOK silencer regions that in turn allows for the expression of a CD8 program in thymocytes^28^.

We and others previously reported an important role for the CD4 and CD8 T cell lineage-defining TFs, ThPOK and Runx3 respectively, in IEL differentiation and Treg function^12, 16, 29, 30 31, 32^ (**Supplementary Fig. 3a**). We hypothesized that ThPOK and Runx3 respective modulation could setinitiate the transcriptional and chromatin changes that ultimately lead to CD4^+^ T cell differentiation towards the IEL fate. We thus asked whether disrupting ThPOK and Runx3 could affect the Treg to IEL progression. To simultaneously fate-map Tregs and abrogate their ThPOK and/or Runx3 expression, we crossed i*Foxp3*^Tomato^ mice with ThPOK^fl/fl^ and/or *Runx3*^fl/fl^ mice^33, 34^ and administered tamoxifen to the i*Foxp3*^Tomato^(ΔThPOK, ΔRunx3, and ΔThPOK ΔRunx3 vs WT) mice over 10 weeks. As previously suggested by our study using a similar strategy^9^, removal of ThPOK promoted the destabilization of Tregs and their acquisition of the a CD4-IEL phenotype, as determined by CD8α expression by Treg–fate-mapped cells, suggesting that the forced deletion of ThPOK leads to premature differentiation of IELs (**Supplementary Fig. 3b, c**). Concomitant ThPOK and *Runx3* ablation in part abrogated the effect of ThPOK deletion, confirming a positive role of Runx3 in CD4-IEL conversion^12^. *Runx3* ablation in Tregs alone did not play a significant role in Treg stability or CD8α-acquisition, implying that it acts primarily in the absence of ThPOK at this stage (**Supplementary Fig. 3b, c**). In contrast to cells in the epithelium, all mLN Tregs remained stable, highlighting the finding that Treg plasticity is tissue–dependent (**Supplementary Fig. 3b, c**). Furthermore, we did not observe any overt signs of disease, suggesting that the remaining stable Tregs, together with the augmented CD4-IELs, were sufficient to maintain intestinal homeostasis.

One key difference between mLN and IE Tregs is that while the former mostly consists of natural Tregs (nTregs), the latter is predominantly composed of less stable peripherally-–induced Tregs (iTregs)^6, 35^ (**Supplementary Fig. 4a**). We next examined if potential effects by ThPOK and Runx3 on Treg plasticity in our *in* vivo results were masked by differences in nTreg vs iTreg stability. Moreover, to dissect the functional roles of ThPOK and Runx3 in these two types of Tregs, we used the classical model of T cell-transfer colitis in *Rag1*-deficient mice^36, 37, 38^. Mice were co-transferred with naïve CD45.1 CD4^+^ T cells along with peripheral CD45.2 Tom^+^ nTregs or iTregs (neuropilin-1 high or low, respectively) sorted from iFoxp3^Tomato^ (ΔThPOK, ΔRunx3, ΔThPOK ΔRunx3 vs WT) mice (**Supplementary Fig. 4b**). While iTregs lost more Foxp3 expression than nTregs as expected, both nTregs and iTregs were stable in the mLN but less so in the IE compartment (**Supplementary Fig. 4c-d**). We again found that ThPOK abrogation led to an increased frequency of ex-Treg-IELs but without a significant loss of Foxp3-expressing cells incompared comparison to the mice transferred with WT iTregs; a result likely relateddue to an increase in Foxp3-CD8α double positive cells (**Supplementary Fig. 4e**). Moreover, concomitant deletion of *Runx3* and ThPOK did not result in increased accumulation of CD8α-expressing cells. In the presence of ThPOK, Runx3 appeared to be stabilizing to iTregs, reinforcing previously proposed roles for this TF in Tregs^30, 31^. Furthermore, we did not observe significant weight loss or intestinal inflammation as shown by low fecal lipocalin-2 levels (**Supplementary Fig. 4f, g)**. Thus, both remaining Tregs orand Tregs that differentiated into CD4-IELs were able to confer protection against colitis. Overall, while deletion of ThPOK was not sufficient to destabilize nTregs as shown defined by Foxp3 expression, in iTregs it led to the development of CD4-IELs in the IE (**Supplementary Fig. 4d, e**).

Our sequencing data reveled that the Treg and IEL programs begin to turn off and on, respectively, at the pre-IEL stage, a transition marked by ThPOK loss. Likewise, our *in* vivo data showed that ablation of ThPOK at the Treg stage led to premature conversion of Tregs to CD4-IELs. To understand the underlying mechanisms set by ThPOK during this progression, we performed RNA and ATAC sequencing on iTregs and exTreg-IELs after tamoxifen-induced deletion of ThPOK in Tregs from i*Foxp3*^tomato^(ΔThPOK) mice (**Fig. 5a**). In agreement with our *in vivo* data, expression of *Foxp3* and related Treg genes was not affected by ThPOK deletion **(Fig. 5b-d, Supplementary Table 7**). Transcriptional profiling of WT and ΔThPOK mLN Tregs revealed that ThPOK loss anticipated the expression of a number of IEL-related transcripts (*Ccl5, Nkg7, Ctsw)* outside the gut environment (**Fig. 5b, e, Supplementary Table 7, 8**). ThPOK–deficient iTregs within the epithelium displayed increased expression of additional IEL genes (*Ccl5, Cd7, Jaml, Cd244a, Nkg7, Cd160, Gzma, Gzmb)* relative to their WT counterparts (**Fig. 5c, e, Supplementary Table 7).** However, only a few gene expression changes between WT and ΔThPOK exTreg-IELs were detected (**Fig. 5d, Supplementary Table 7**), an expected result given that ThPOK is naturally downregulated in exTreg-IELs. Principle component analysis (PCA) and Euclidean distances comparing the transcriptional profiles of all sequenced WT and ΔThPOK cell types in the IE confirmed that abrogation of ThPOK has the most effect at the iTreg stage. The profiles of both iTregs and ex-Treg-IELs in the IE of ΔThPOK mice more closely resembled the WT pre-IELs than their WT counterparts, indicating that forced ThPOK loss at the Treg stage prematurely pushes cells to differentiation to IELs before the Treg program is shut down (**Fig. 5f, g**). This was also suggested by the increase in Foxp3^+^CD8α^+^ CD4^+^ T cells we observed in the IE of mice transferred with ΔThPOK iTregs.

**Figure 5:**
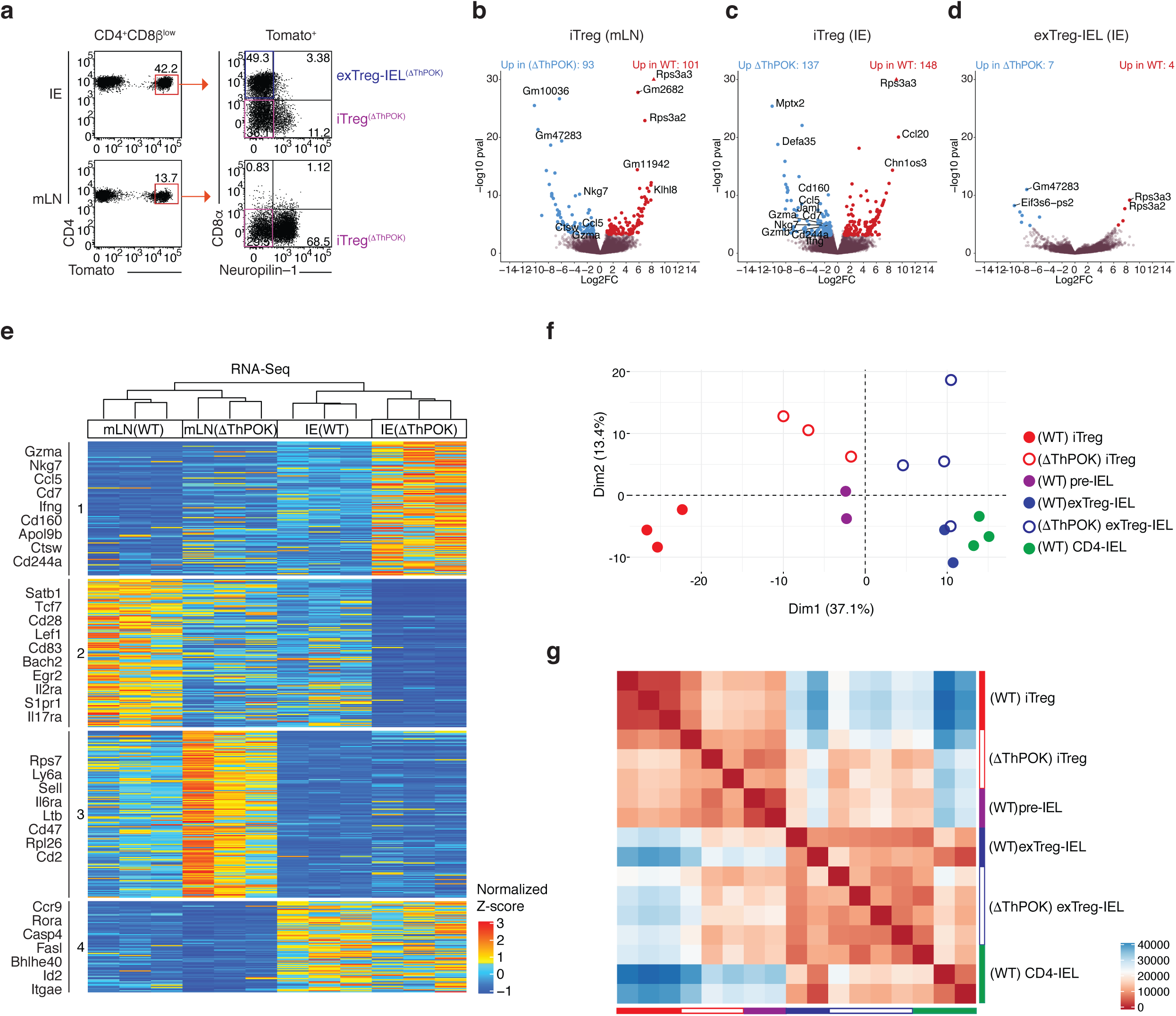
Abrogation of ThPOK in Tregs anticipates progression to IELs at the transcriptional level. (**a-g**) Induced Tregs (iTregs; Tomato^+^CD8α^−^neuropilin-1^−^) and Treg-derived CD4-IELs (exTreg-IELs; Tomato^+^CD8α^+^) were sorted from *Zbtb7b*^fl/fl^x*Runx3*^fl/+^x*Rosa26*^|s|tdTomato^ x*Foxp3*^CreER^ (i*Foxp3*^(Δ*ThPOK*)^) mice after 10 weeks of tamoxifen administration followed by RNA-Sequencing from IE or mLN. (**a**) ΔThPOK T cell populations sequenced. (**b-d**) Volcano representation of differentially expressed genes (DEG) between WT and ΔThPOK of the same cell type as indicated, performed by Wald pairwise comparison test, p_adj_< 0.05. (**e**) Heatmap of DEGs between indicated bulk populations. Expression values represents the normalized Z-score of gene abundances (TPM) (**f, g**) Principle component analysis (**f**) and corresponding Euclidean distance analysis (**g**) of DEG of all WT and ΔThPOK cell types from the IE. (n=3-4 i*Foxp3*^(Δ*ThPOK*)^). Each sample consisted of 300-800 cells per mouse (n=2-4 mice).

Pairwise comparisons of accessible chromatin between WT and ΔThPOK counterparts revealed minimal effects of ThPOK ablation at the Treg stage, corroborating our *in vivo* data and suggesting that the effects of ThPOK loss primarily result in transcriptional changes relative to chromatin accessibility (**Fig.6a, b, Supplementary Table 9**). About 30-40% of all differences in chromatin accessibility between the genotypes occur at promoter regions, a proportion in line with that of ThPOK binding sites (**Fig. 6c**). Similar to the RNA-Seq analysis, PCA and corresponding Euclidean distances of the ATAC-Seq data also revealed that the ablation of ThPOK at the Treg stage results in both iTregs and exTreg-IELs from ΔThPOK mice to more closely resemble WT pre-IELs (**Fig. 6d, e**). Taken together, our RNA and ATAC-sequencing data show that the untimely downmodulation of ThPOK at the Treg stage results in increased chromatin accessibility and transcription of the IEL program prior to the shutting down of the Treg program, resulting in IELs that resemble destabilized Tregs at the pre-IEL stage. This suggests that the natural IE–induced downmodulation of ThPOK allows for the full loss of the Treg program and acquisition of the IEL program in succession. Taken together, these analyses suggest that while ThPOK downmodulation initiate the acquisition of an IEL program already outside the gut, the epithelial environment is still critical for full IEL differentiation.

**Figure 6:**
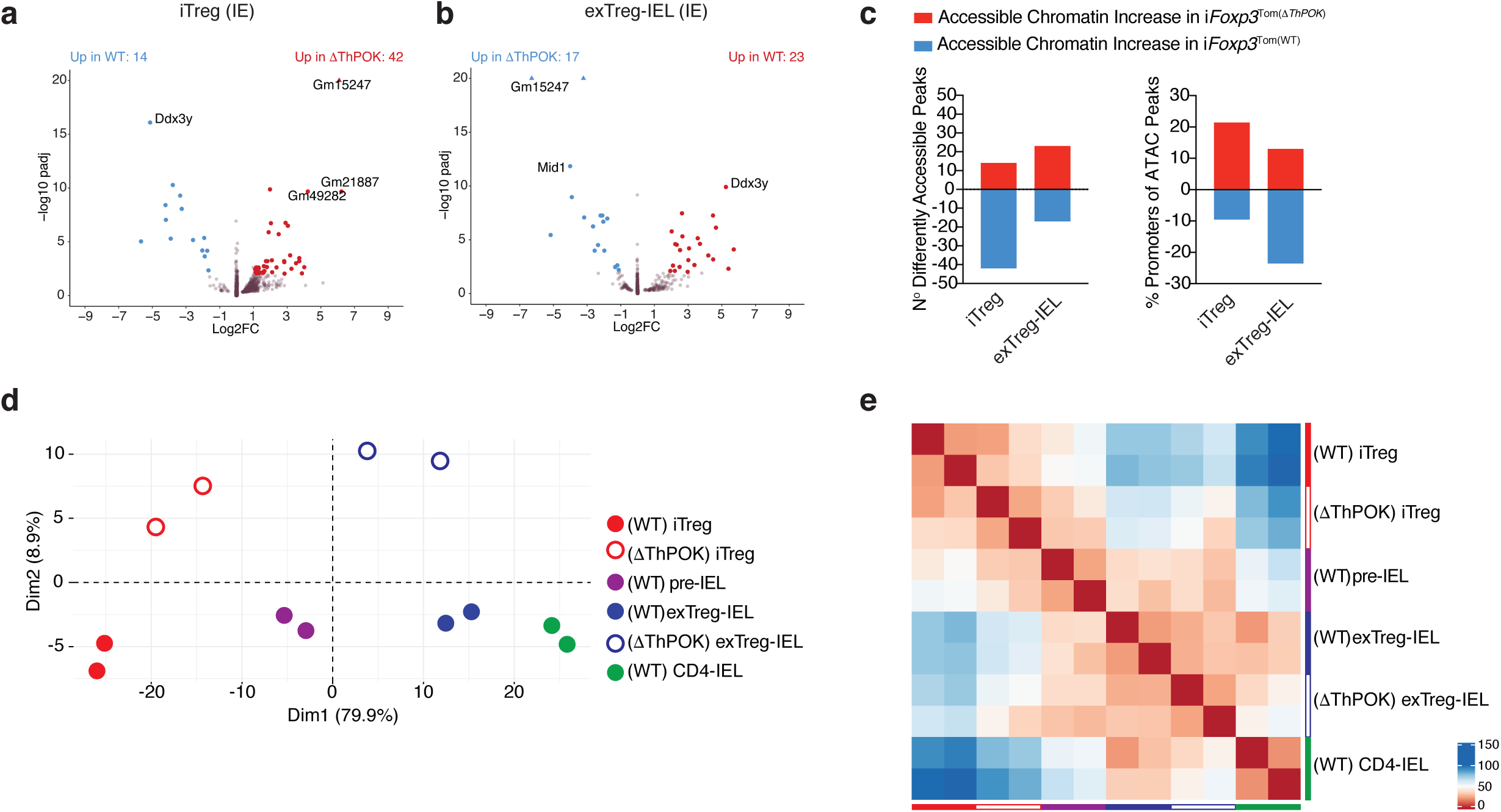
Abrogation of ThPOK in Tregs enhances progression to IELs at the chromatin level. (**a-e**) Induced Tregs (iTregs; Tomato^+^CD8α^−^neuropilin-1^−^) and Treg derived CD4-IELs (exTreg-IELs; Tomato^+^CD8α^+^) were sorted from *Zbtb7b*^fl/fl^x*Runx3*^fl/+^x*Rosa26*^|s|tdTomato^ x*Foxp3*^CreER^ (i*Foxp3*^(Δ*ThPOK*)^) mice after 10 weeks of tamoxifen administration followed by ATAC-Sequencing from IE. (**a, b**) Volcano representation of differentially accessible chromatin regions (DACR) between WT and ΔThPOK of the same cell type as indicated, performed by Wald pairwise comparison test, p_adj_<0.01. (**c**) Data of accessible chromatin increase (red) and decrease (blue) between WT and ΔThPOK of the same cell type as indicated. Numbers of DACR (left) and percent of DACRs at promoter regions (right). (**d-e**) Principle component analysis (**d**) and corresponding Euclidean distance heatmap (**e**) of DACR of all WT and ΔThPOK cell types from the IE. Each sample consisted of 5,000-40,000 cells from 2 or 3 pooled mice (n=2 samples).

## Discussion

In this study, we coupled genetic fate-mapping and gene ablation mouse models with RNA and ATAC sequencing to elucidate how the intestinal tissue induces T cell plasticity from the gut–draining LNs to the intestinal wall. Our unbiased survey of transcriptional profiles of CD4^+^ T cells in three enteric–associated sites revealed a strong inter– and intra–tissue adaptation, which included chromatin and transcriptional changes towards an IEL program within the gut epithelium.

We uncovered a stepwise process imprinted on T cells upon migrating to the epithelium, regardless of the subset of origin. Despite a very similar transcriptional profile between IELs derived from Tregs and conventional CD4^+^ T cells, exTreg-IELs retained low levels of the chromatin profile from their Treg precursors. This finding corroborates a previous study showing that changes in chromatin accessibility and transcription did not always match as Tregs adapted to different tissues^39^. Nevertheless, our identification of a pre-IEL signature, marked by the loss of key features of conventional CD4 and/or Treg programs, suggests that during tissue adaptation a cell must first shut down transcriptional programs in place (or decouple TFs bound to its DNA targets) that may prevent tissue imprinting, suggestion corroborated by previous studies describing roles of Runx3 to the CD8-lineage or tissue–resident memory T cells (T_RM_)^26, 27^. A recent study also describes a stepwise process taken by Tregs, in which peripheral precursors gradually acquire chromatin accessibility and reprograming towards nonlymphoid–tissue Tregs. The authors identified the transcription factor BATF as a main driver of the molecular tissue programing in the precursors^40^. These observations parallel our findings regarding epithelial imprinting on Tregs; while Runx3 and ThPOK seemed dispensable in Treg function in the transfer colitis model, ThPOK downmodulation was a driver of induced Treg plasticity towards the IEL fate at the epithelium. It remains to be determined, like the BATF–differentiation described, whether lamina propria or epithelium Tregs share a peripheral common precursor prone to tissue differentiation in the gut.

Our previous studies pointed to an important role for ThPOK in Treg stability as assessed by the continuous expression of Foxp3 despite ThPOK ablation^9^. If so, prolonged ThPOK inactivation in Tregs should result in overt autoimmunity. Our *in vivo* experiments presented here seem to refute this possibility, as Treg suppressive function appeared unaltered, while Foxp3 expression was only modestly affected. Similarly, while *Thpok* deletion in activated CD4^+^ T cells does not affect viral clearance or the expansion of antigen-specific cells following LCMV infection, it plays a role in the regulation of CD4^+^ T cell differentiation, particularly towards Th1 cells, which share some expression profile with interferon-producing CD4-IELs^24^. One caveat of broad conclusions regarding the role of ThPOK in Treg programming is that our sequencing analyses revealed that *Foxp3* is one of the last canonical Treg genes to be downmodulated during the IEL transition, which reinforces previous studies suggesting that Foxp3 is a late-acting transcription factor which binds to pre-established accessible enhancer regions^41^. Therefore, it is plausible that the Treg program may be modulated independently of Foxp3; in the case of tissue adaptation events addressed here, this may preclude the use of Foxp3 as Treg marker without considering the entire Treg profile. Additionally, it is possible that ThPOK acts in conjunction with other TFs, including Runx3^30, 31, 32^, to regulate Treg function. By inactivating both ThPOK and its homolog LRF in differentiated Tregs, a previous study concluded that these transcription factors redundantly support Foxp3 function^29^. Finally, the differential methylation patterns in the regulatory regions on the *Foxp3* locus between natural and induced Tregs^42^ raises the possibility that they display distinct dependence on ThPOK, as the latter are more prone to losing Foxp3 expression both *in vivo* and upon transfer to lymphopenic environments. Although our work elucidated the events following ThPOK downmodulation in Tregs, the exact tissue–dependent triggers leading to ThPOK downmodulation is not completely known^12, 13, 16^. A microbiota induced aryl-hydrocarbon receptor requirement of this crucial step has been proposed to drive ThPOK downmodulation, and it would be interesting to understand this and additional mechanisms in greater detail^43^.

Lymphocytes are thought to display common features inherent to the tissue in which they reside ^21, 39, 44, 45, 46^. Consequently, migrating immune cells must be able to readily adapt to their new environment at the transcriptional level, although the type and speed of tissue-imprinting will vary with the tissue and the immune cell of interest^20, 39, 45, 46, 47, 48, 49, 50^. For instance, it was proposed that while Tregs from skin and colon retain a common Treg program, they concomitantly acquire tissue-specific profiles^21^. Conversely, studies characterizing T_RM_, which share several characteristics with IEL subsets, also observed a progressive acquisition of a tissue–specific transcriptional profile, distinct of circulating memory cells^46^. Our studies add to this line of investigation by defining a relationship between two distinct but adjacent intestinal compartments and their draining LNs; our findings indicate that peripheral T cells are likely to either go to the intestinal lamina propria and reside there as distinct CD4^+^ subsets or proceed to the epithelial compartment without being imprinted with a LP signature but become polarized with an IEL profile. These observations point to important distinctions between these two closely juxtaposed intestinal layers and may call for specific approaches to target cells located in each site. Additionally, they raise the possibility that similar reciprocal trajectories take place in other sites, such as the skin, between dermal and epidermal compartments^48^, or the lung, between the stroma and epithelium^49, 50^. We conclude that T cells undergo wide, tissue–specific and stepwise chromatin and transcriptional changes from lymphoid to non-lymphoid tissue sites, highlighting the existence of a late stage of lymphocyte plasticity likely required for the maintenance of tissue homeostasis.

## Methods

### Animals

Animal care and experimentation were consistent with the NIH guidelines and were approved by the Institutional Animal Care and Use Committee at the Rockefeller University. *Rag1*^-/-^ (002216), *Zbtb7b*^eGFP^(027663), *Zbtb7b*^*fl/fl* 33^(009369), *Rosa26*^*|s|tdTomato*^(007914), *Foxp3*^eGFP-Cre-ERT2^(016961), *Foxp3*^IRES-mRFP^(008374), *Cd4*^Cre-ERT2^(022356) and B6.SJL-*Ptprc*^a^*Pepc*^b^(002014) mice were purchased from the Jackson Laboratories and maintained in our facilities. *Runx3*^fl/f*l*^ *(008773)* mice were kindly provided by T. Egawa (Wash. U.). Several of these lines were interbred in our facility to obtain the final strains described in the text. Genotyping was performed according to the protocols established for the respective strains by Jackson Laboratories. Mice were maintained at the Rockefeller University animal facilities under specific pathogen-free (SPF) conditions.

### Antibodies and flow cytometry analysis

Fluorescent dye–conjugated antibodies were purchased from BD Biosciences, Biolegend or Ebioscience (Thermofisher). The following clones were used: anti-CD45.1, A20; anti-Foxp3, FJK-16s; anti-CD4, RM4-5; anti-CD45.2, 104; anti-CD8α, 53-6.7; anti-CD8β, YTS 156.7.7; anti-CD44, IM7; anti-CD45, 30-F11; anti-CD62L, MEL-14; G8.8; anti-TCRβ, H57-597; anti-TCRγδ, eBioG23; anti-CD25, PC61.5;. Live/dead fixable dye Aqua (ThermoFisher Scientific) was used according to manufacturer’s instructions. Intracellular staining of Foxp3 was conducted using Foxp3 Mouse Regulatory T Cell Staining Kit (eBioscience, USA). Flow cytometry data was acquired on a LSR-II flow cytometer (Becton Dickinson, USA) and analyzed using FlowJo software package (Tri-Star, USA). Anti-ThPOK ChIP antibody was kindly provided by T. Egawa (Wash. U.).

### Isolation of intestinal T cells

Intraepithelial and lamina propria lymphocytes were isolated as previously described ^12, 15^. Briefly, small intestines were harvested and washed in PBS and 1mM dithiothreitol (DTT) followed by 30 mM EDTA. Intraepithelial cells were recovered from the supernatant of DTT and EDTA washes and mononuclear cells were isolated by gradient centrifugation using Percoll. Lamina propria lymphocytes were obtained after collagenase digestion of the tissue. Single-cell suspensions were then stained with fluorescently labeled antibodies for 20min at 4°C prior to downstream flow cytometry (analysis or sorting) as specified in figure legends.

### Tamoxifen treatment

For *in vivo* treatment, mice were intragastrically administered with 5 mg of tamoxifen (Sigma) dissolved in corn oil (Sigma) and 10% ethanol at 50 mg/ml. Tamoxifen was administered to mice starting at 6-7 weeks old, 4 times in the first week and then 2 times every week (3 days apart) every other week for 8-10 weeks. To label cells for transfer, tamoxifen was administered only twice, 2 days apart, 2 days prior to sorting. 2 days after transfer to *Rag1*^-/-^ mice, animals were administered 100uL of 10mg/mL Tamoxifen intraperitonially.

### Cell sorting

Lymphocytes were sorted on a FACS Aria II instrument as indicated in the figure legends.

### Cell transfer

Induced (neuropilin-1^−^) and natural (neuropilin-1^+^) Tregs (CD45.2^+^TCRβ^+^CD4^+^CD8^−^CD25^+^Tomato^+^) from spleen and mLN of donor CD45.2 mice were sorted 2 days after mice were intragastrically given 5mg of tamoxifen prepared as described above for 2 days and co-transferred with naïve CD4^+^ T cells (CD45.1^+^TCRβ^+^CD4^+^CD8^-^CD62L^high^CD44^low^) sorted from CD45.1 donor mice and 100,000 CD45.2^+^ Tregs with 400,000 CD45.1^+^ naïve T cells were intravenously transferred to *Rag1*^-/-^ hosts. Body weight and fecal lipocalin-2 levels were monitored until terminal analysis.

### ELISA for Lipocalin-2

Lipocalin-2 was analyzed by using Lcn-2 ELISA kit (R&D, MN) as described^51^.

### ATAC-Sequencing

ATAC-Seq was performed as previously described^22, 52^ on 5,000-40,000 FACS-purified cells from 2-9 mice. In brief, cells were lysed in lysis buffer for 1 minute and transposed with Tagment DNA Enzyme 1 (Illumina) for 30 minutes. DNA was cleaned up using a MinElute DNA purification Kit (Qiagen), followed by barcoding and library preparation by the Nextera DNA Library preparation kit (Illumina) according to manufacturer’s guidelines and sequenced on an Illumina NextSeq500.

### ChIP-Sequencing

Cells were fixed in 1% formaldehyde for 20min, quenched with 0.15M glycine and washed in PBS. 2×10^6^ were then sorted and lysed for 30min at 4°C. Cells were then sonicated for 22 minutes at 30 seconds on/off using the Bioruptor sonicator (Diagenode) and spun down. 10% was frozen for input, the remaining 90% was incubated with anti-ThPOK antibody bound to goat anti-rabbit M280 magnetic beads (Invitrogen) overnight at 4°C prior to washing with RIPA buffer and overnight decrosslinking at 65°C. DNA was then eluted off beads into TE buffer and, along with input, purified using the Zymogen DNA clean & concentrator kit. DNA was sequenced using NextSeq2500. To obtain Tregs, Foxp3^RFP^ reporter mice were injected with IL-2/αIL-2 complex as previously described for 3 consecutive days for Treg expansion^53, 54^. CD4^+^ T cells were isolated from the spleen and mLNs and sorted as Thy1^+^TCRβ^+^CD4^+^CD8α^−^RFP^+^.

### Bulk RNA-Sequencing

Sorted cells (300-800) were lysed in TCL buffer (Qiagen, 1031576) supplemented with 1% β-mercaptoethanol. RNA was isolated using RNAClean XP beads (Agentcourt, A63987), reversibly transcribed, and amplified as described^55^. Uniquely barcoded libraries were prepared using Nextera XT kit (Illumina) following manufacturer’s instructions. Sequencing was performed on an Illumina NextSeq550.

### Single cell RNA-Seq library preparation

Lymphocytes were sorted, counted for viability and immediately subjected to library preparation. The scRNA-Seq library was prepared using the 10x Single Cell Chromium system, according to the manufacturer’s instructions at the Genomics core of Rockefeller University and was sequenced on an Illumina NextSeq550 to a minimum sequencing depth of 50,000 reads per cell using read lengths of 26 bp read 1, 8 bp i7 index, 98 bp read 2.

### Statistical analysis

Statistical analysis was performed using GraphPad Prism software. Data was analyzed by applying Student’s *t* test, Mann-Whitney test, one-way ANOVA with Tukey post-test, Log-rank test or Kruskal-Wallis with Dunns post-test whenever necessary. For analysis of non-parametric histological scores, a Mann-Whitney test was used. A *P* value of less than 0.05 was considered significant. * *P*<0.05, ** *P*<0.01, *** *P*<0.001.

### Single Cell RNA-Seq analysis

The single cell RNA-Seq fastq files originated from the Chromium 10x libraries were processed with Cell Ranger (v. 3.0.2) for cell barcode aggregation, genome alignment, and UMI transcript quantification. The matrix of gene counts was analyzed with the Seurat (v. 3.1.2) package for the R environment ^56^. For downstream analysis, the top 3000 genes containing the highest standardized variance and expression were pre-selected. The detected cell clusters were visualized by performing Uniform Manifold Approximation and Projection (UMAP) algorithm ^57^. Cluster signatures were determined by the Wilcoxon Rank Sum test and genes containing adjusted p-values smaller than 0.01 were considered significant. For pseudotime trajectory analysis, the Seurat dataset was converted into a ‘SingleCellExperiment’ object and used within Slingshot and Monocle3^58^. Both Slingshot and Monocle3 algorithm was performed by setting the cluster 10 (naïve cells from mLN) as the root.

### Bulk RNA-Seq data analysis

The raw fastq sequencing files were processed together with the gencode mouse annotation database (v. M21), by running kallisto (v. 0.46.0) to calculate the transcript abundances^59^. Next, the abundance files were submitted to the sleuth (v. 0.30.0) pipeline based on R, for transcript abundance normalization, gene expression analysis and statistics^60, 61^. The quantified transcripts were combined into genes for the downstream analysis. Batch effects were evaluated through principal component analysis and removed with Limma^62^. To capture significantly expressed genes we performed a likelihood ratio test between a null model and our experimental design using an adjusted p-value of 0.05 as threshold. The selected features were clustered using k-means and represented as a heatmap to define group gene signatures. Pairwise comparisons between groups were performed using the Wald-test and significant genes were considered for downstream analysis by having an adjusted p-value smaller than 0.05 and a log2 fold-change of 1. Pre-ranked gene set enrichment analysis (GSEA) were performed with the fgsea package^63^ by comparing gene lists sorted by their log2 fold-change with gene signatures from the clusters 21 (Activated-Treg) and 6 (CD4-IEL) of the single cell dataset (10x Genomics).

### ATAC-Seq analysis

The raw bulk ATAC-Seq files were processed using the ENCODE-DCC pipeline (https://github.com/ENCODE-DCC/atac-seq-pipeline) automated through Cromwell (https://github.com/broadinstitute/cromwell). Shortly, adapter sequences were trimmed (Cutadapt) prior to genome mapping (Bowtie2)^64^ and filtered subsequently filtered (Samtools)^65^. Next, sequencing enriched regions, associated to chromatin accessible peaks were called (MACS2) and filtered (IDR) by running Irreproducible Discovery Rate^66^ algorithm with a 10% threshold, meaning 10% chance of being an irreproducible peak. Later, filtered peak regions were annotated by using HOMER software tools. Comprehensive motif screening was performed with MEME-ChIP^25^ by using promoter originated peaks and, length-normalized reads to a 500 kb to a fixed window of 500bp. Enriched transcriptional factor motifs on ATAC-Seq peaks were detected with Centrimo^67^ by providing known Human Runx3 (MA0684) and Foxp3 (MA0850) as published in JASPAR 2020 ^68, 69^. As indicative of the presence of Zbtb7b (Thpok) we used the enriched motif detected in our Thpok ChIP-Seq peaks.

### ChIP-Seq analysis

The raw bulk ChIP-Seq files were processed using the ENCODE-DCC pipeline (https://github.com/ENCODE-DCC/chip-seq-pipeline2) automated through Cromwell (https://github.com/broadinstitute/cromwell). Shortly, adapter sequences were trimmed (Cutadapt) prior to genome mapping (BWA) and filtered subsequently (Samtools)^65, 70^. Next, sequencing enriched regions, associated to chromatin accessible peaks were called (SPP) and filtered (IDR) by running Irreproducible Discovery Rate algorithm with a 5% threshold, meaning 5% chance of being an irreproducible peak^71, 72^. Later, filtered peak regions were annotated by the HOMER software suite of tools^73^. Comprehensive motif screening was performed with MEME-ChIP by using promoter originated, length-normalized reads to a 500 kb.

## Acknowledgements

We are grateful to A. Rogoz, and S. Gonzalez for exceptional animal care, mouse colony management and genotyping and the Rockefeller University employees for continuous assistance. We thank K. Gordon and K. Chhosphel for assistance with cell sorting. We thank C. Zhao and the entire Genomics Core of Rockefeller University for library preparation for 10X Genomics and assistance with all sequencing platforms used in this paper. We are grateful to U. Schaefer (The Rockefeller University) for help with ChIP protocols and guidance, S. Larsen for help with the ATAC-Seq and T. Sujino (Keio University) for continuous discussion and help with mouse breeding. We thank B. Reis and G. Victora for suggestions and critical reading of the manuscript, and all the members of the Mucida lab for fruitful discussions.

## Author contribution

DM conceived the study. ML, AMB and DM designed experiments and wrote the manuscript. ML and AMB performed experiments. TBRC performed all bioinformatics analyses. TBRC, ML and AMB analyzed experiments.

## Figure Legends

**Supplementary Figure 1.**
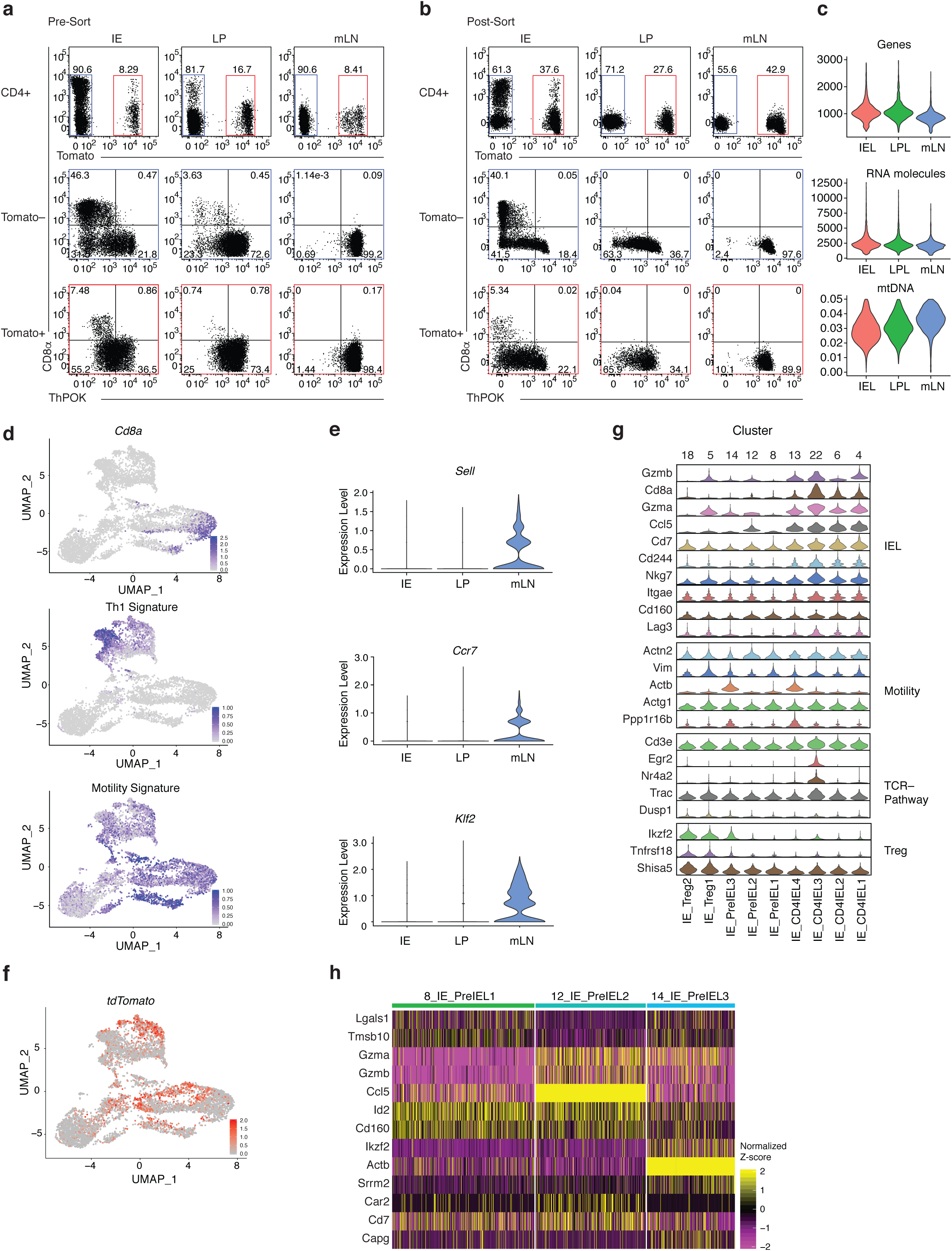
Related to Figure 1. (**a-h**) i*Foxp3*^Tom^ThPOK^GFP^ mice were treated with tamoxifen for 10 weeks, and Tomato^−^ and Tomato^+^ CD4^+^ T cells from mesenteric lymph nodes (mLN), lamina propria (LP) and intestinal epithelium (IE) were sorted for scRNA-Seq using 10X Genomics platform. Sorted Tomato^−^ (blue gates) and Tomato^+^ (red gates) cells were pooled in a 2:1 ratio per tissue, resulting in 3 separate libraries. (**a, b**) CD4^+^ T cells from mLN, LP and IE before sorting (**a**) and after sorting (**b**). (**c**) Number of sequenced genes (top) and RNA molecules (middle) per cluster and percent of mitochondrial DNA (bottom) per library. (**d**) Expression levels of *Cd8a* (top), the Th1 signature (middle) and the motility signature (bottom) of all sequenced cells. (**e**) Expression levels of *Sell* (top), *Ccr7* (middle), and *Klf2* (bottom) of cells from indicated tissues. (**f**) *tdTomato* gene expression in 6668 sequenced cells. (**g**) Expression levels of indicated genes among IE clusters. (**h)** Expression heatmap of selected genes in pre-IEL clusters.

**Supplementary Figure 2:**
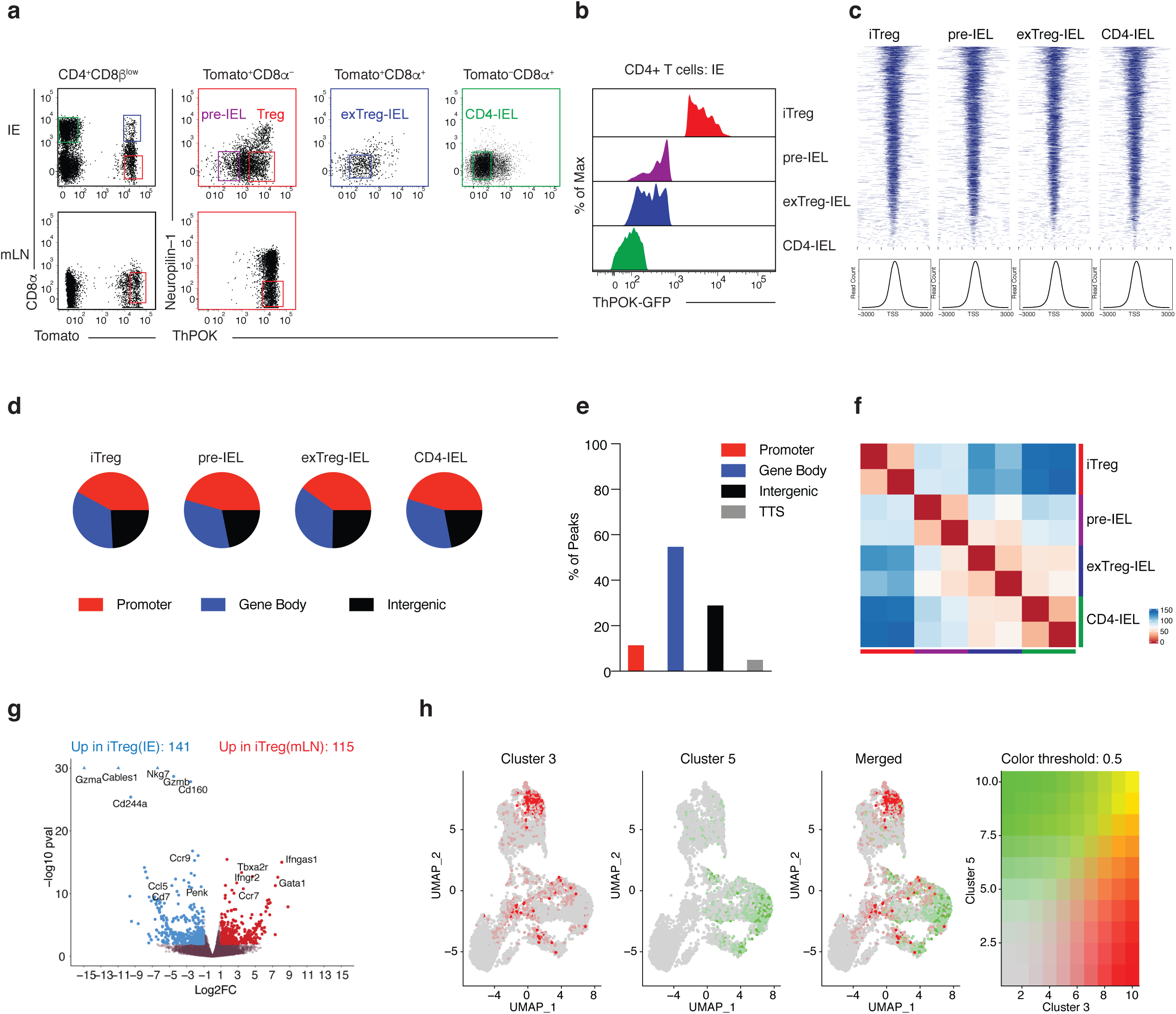
Related to Figure 3. (**a-h**) i*Foxp3*^Tom^ThPOK^GFP^ mice were treated with tamoxifen for 10 weeks and induced Tregs (iTreg; CD4^+^Tomato^+^GFP^High^neuropilin-1^−^CD8α^−^), pre-IELs (CD4^+^Tomato^+^ GFP^Low^CD8α^−^), exTreg-IELs (CD4^+^Tomato^+^GFP^Low^CD8α^+^), and CD4-IELs (CD4^+^ Tomato^−^GFP^Low^CD8α^+^) were sorted in bulk from the IE. Assay for transposase-accessible chromatin (ATAC) or RNA libraries were prepared followed by sequencing of indicated populations. iTregs were also sorted from the mLN for RNA-seq. (**a**) Sorting strategy of indicated populations. CD8α and Tomato expression among CD4^+^ CD8β^low^ T cells (left) from IE (top) and mLN (bottom). ThPOK and neuropilin-1 expression among Tomato^+^CD8α^−^ (red), Tomato^+^CD8α^+^ (purple) and Tomato^−^CD8α^+^ (green) cells from IE (top) and mLN (bottom). (**b**) Levels of ThPOK expression in each sorted population in the IE. (**c**) Mapped accessible chromatin regions relative to transcriptional start sites (TSS) of all genes in ATAC-Seq data (top) and their relative read counts per genomic regions (bottom). (**d**) ATAC-Seq peak annotations as indicated per cell type. (**e**) Percent of total differentially accessible chromatin regions as follows: 5’UTR and promoters (Promoter; red), 3’UTR with exons and introns (Gene Body; blue), transcriptional termination site (TTS; grey) and intergenic (black). (**f**) Euclidean distance correlation of chromatin accessibility profiles of all samples. (**g**) Volcano representation of differentially expressed genes between IE iTregs (higher expression in blue) and mLN iTregs (higher expression in red), performed by Wald pairwise comparison test, p_adj_<0.05 values were considered significant. (**h**) Treg signature from cluster 3 and IEL signature from cluster 5 of the bulk RNA-seq heatmap (Figure 3d) overlaid onto the scRNA-Seq UMAP from figure 1.

**Supplementary Figure 3.**
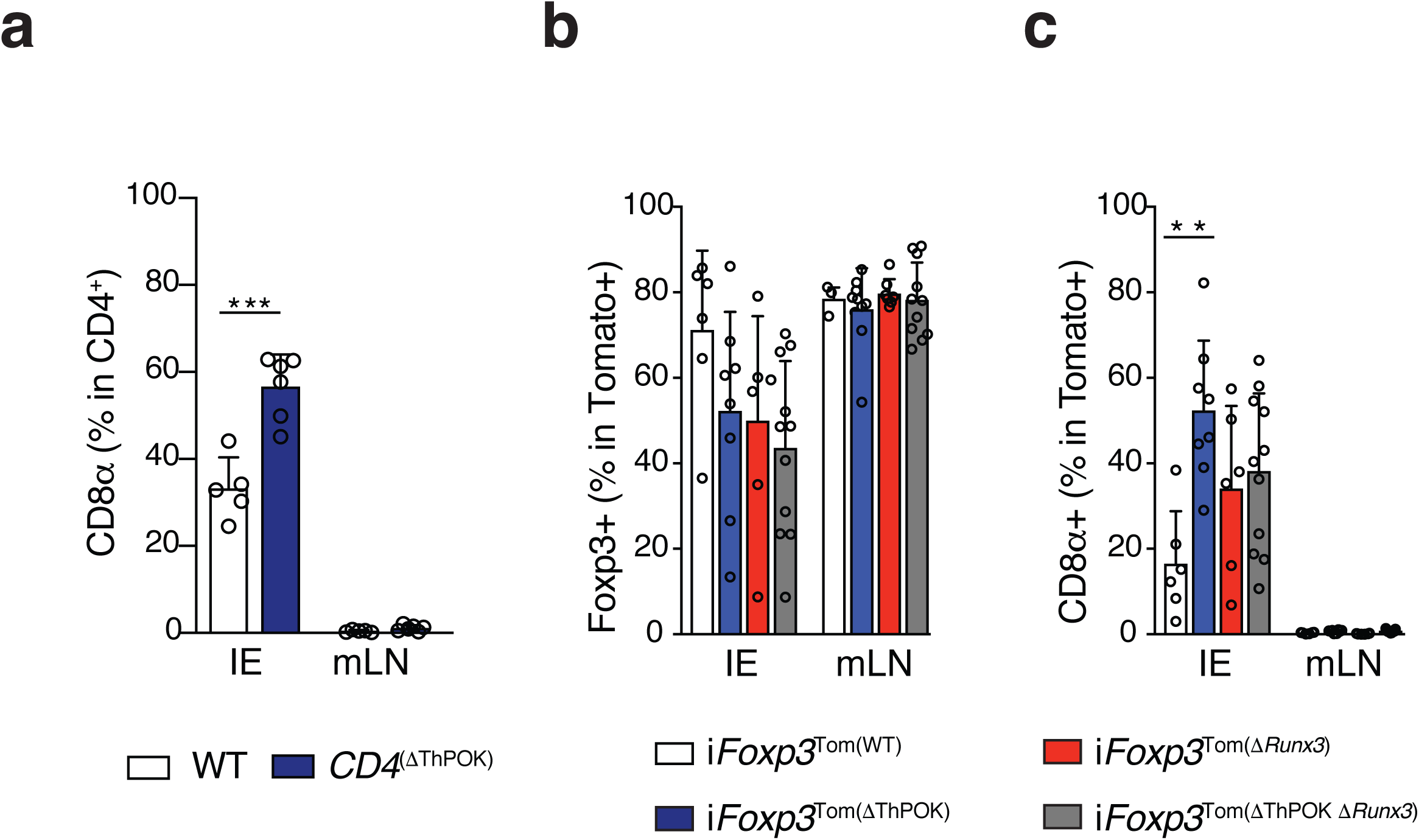
ThPOK downmodulation leads to CD8α induction in intraepithelial Tregs. (**a**) Flow cytometry analysis of CD8α^+^ among CD4^+^ T cells from the intraepithelial compartment (IE) and mesenteric lymph nodes (mLN) in *Cd4*^CreERT2–^x*Zbtb7b*^fl/fl^ (WT) or *Cd4*^CreERT2+^x*Zbtb7b*^fl/fl^ (*Cd4*^(Δ*ThPOK*)^) animals after 10 weeks of tamoxifen treatment. (**b, c**) Flow cytometry analysis of CD45^+^TCRβ^+^CD4^+^CD8b^low^ Tomato^+^ cells in the IE and mLN of *Zbtb7b*^fl/+^x*Runx3*^fl/+^x*Rosa26*^|s|tdTomato^x*Foxp3*^CreER^ (i*Foxp3*), i*Foxp3*x*Runx3*^fl/+^ (i*Foxp3*^(Δ*ThPOK*)^), i*Foxp3*x*Zbtb7b*^fl/+^x*Runx3*^fl/fl^ (i*Foxp3*^(Δ*Runx3*)^), i*Foxp3*x*Zbtb7b*^fl/fl^x*Runx3*^fl/fl^ (i*Foxp3*^(Δ*ThPOK* Δ*Runx3*)^) mice after 10 weeks of tamoxifen treatment. (**b**) Frequency of total Foxp3^+^ cells among Tomato^+^ CD4^+^ T cells. (**c**) Frequency of total CD8α^+^ cells among Tomato^+^ CD4^+^ T cells. Data are expressed as mean +/- SEM of individual mice (n=5-11 per genotype, 3 separate experiments). *p<0.05, **p<0.01, ***p<0.001 (one-way ANOVA and Bonferonni test).

**Supplementary Figure 4.**
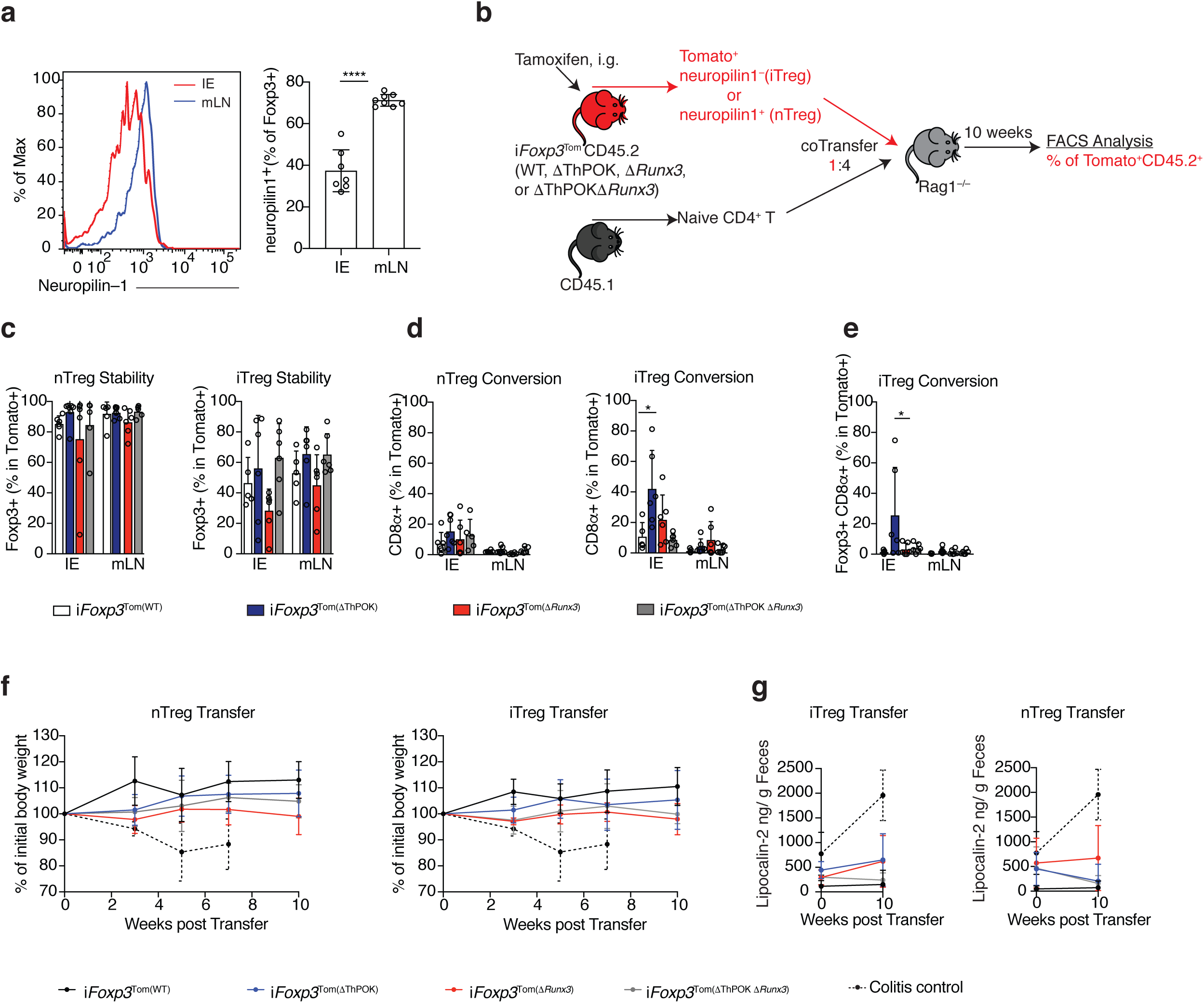
Impact of ThPOK and Runx3 on Treg stability and function. (**a**) Neuropilin-1 expression among Foxp3^+^ Tregs in the intraepithelial compartment (IE) and mesenteric lymph nodes (mLN) of WT mice. Histogram (left) and frequency (right). (**b-g**) nTreg (neuropilin-1^−^) or iTregs (neuropilin-1^+^) were sorted from spleens and mLNs of CD45.2 *Zbtb7b*^fl/+^x*Runx3*^fl/+^x*Rosa26*^|s|tdTomato^x*Foxp3*^CreER^ (i*Foxp3*), i*Foxp3*x*Runx3*^fl/+^ (i*Foxp3*^(Δ*ThPOK*)^), i*Foxp3*x*Zbtb7b*^fl/+^x*Runx3*^fl/fl^ (i*Foxp3*^(Δ*Runx3*)^), i*Foxp3*x*Zbtb7b*^fl/fl^x*Runx3*^fl/fl^ (i*Foxp3*^(Δ*ThPOK* Δ*Runx3*)^) mice after tamoxifen administration and co-transferred with CD45.1 naïve CD4^+^ T cells to *Rag1*^-/-^ hosts. CD45.2^+^TCRβ^+^CD4^+^CD8β^-^Tomato+ lymphocytes from the IE and mLN were analyzed 10 weeks after transfer. (**b**) Experimental layout. (**c, d**) Frequencies of total intracellular Foxp3 (**c**) and total surface CD8α (**d**) cells after nTreg (left) or iTreg (right) transfer. (**e**) Frequency of Foxp3^+^CD8α^+^ cells after iTreg transfer. (**f-g**) Body weight of *Rag1*^-/-^ recipients after nTreg (left) or iTreg (right) transfers (**f**) and levels of fecal lipocalin-*2* before and after per cell type transferred (**g**). Dashed lines represent colitis control transfer of naïve CD45.1^+^ CD4^+^ T cells only. Data are expressed as mean +/- SEM of individual mice (n=5-6 per genotype, 5 experiments). *p<0.05, **p<0.01, ***p<0.001 (one-way ANOVA and Bonferonni test).

